# The Role of State Uncertainty in the Dynamics of Dopamine

**DOI:** 10.1101/805366

**Authors:** John G. Mikhael, HyungGoo R. Kim, Naoshige Uchida, Samuel J. Gershman

**Author notes:** These authors contributed equally to this work (John G. Mikhael, HyungGoo R. Kim).

## Abstract

Reinforcement learning models of the basal ganglia map the phasic dopamine signal to reward prediction errors (RPEs). Conventional models assert that, when a stimulus predicts a reward with fixed delay, dopamine activity during the delay should converge to baseline through learning. However, recent studies have found that dopamine ramps up before reward in certain conditions even after learning, thus challenging the conventional models. In this work, we show that sensory feedback causes an unbiased learner to produce RPE ramps. Our model predicts that, when feedback gradually decreases during a trial, dopamine activity should resemble a ‘bump,’ whose ramp-up phase should furthermore be greater than that of conditions where the feedback stays high. We trained mice on a virtual navigation task with varying brightness, and both predictions were empirically observed. In sum, our theoretical and experimental results reconcile the seemingly conflicting data on dopamine behaviors under the RPE hypothesis.

## Introduction

Perhaps the most successful convergence of reinforcement learning theory with neuroscience has been the insight that the phasic activity of midbrain dopamine (DA) neurons tracks ‘reward prediction errors’ (RPEs), or the difference between received and expected reward (Schultz et al., 1997; Schultz, 2007a; Glimcher, 2011). In reinforcement learning algorithms, RPEs serve as teaching signals that update an agent’s estimate of rewards until those rewards are well-predicted. In a seminal experiment, Schultz et al. (1997) recorded from midbrain DA neurons in primates and found that the neurons responded with a burst of activity when an unexpected reward was delivered. However, if a reward-predicting cue was available, the DA neurons eventually stopped responding to the (now expected) reward and instead began to respond to the cue, much like an RPE (see Results). This finding formed the basis for the RPE hypothesis of DA.

Over the past two decades, a large and compelling body of work has supported the view that phasic DA functions as a teaching signal (Schultz et al., 1997; Niv and Schoenbaum, 2008; Glimcher, 2011; Steinberg et al., 2013; Eshel et al., 2015). In particular, phasic DA activity has been shown to track the RPE term of temporal difference (TD) learning models, which we review below, remarkably well (Schultz, 2007a). However, recent results have called this model of DA into question. Using fast-scan cyclic voltammetry in rat striatum during a goal-directed spatial navigation task, Howe et al. (2013) observed a ramping phenomenon— a steady increase in DA over the course of a single trial—that persisted even after extensive training. Since then, DA ramping has been observed during a two-armed bandit task (Hamid et al., 2016), during the execution of self-initiated action sequences (Collins et al., 2016), and in the timing of movement initiation (Hamilos et al., 2020). At first glance, these findings appear to contradict the RPE hypothesis of DA. Indeed, why would error signals persist (and ramp) after a task has been well-learned? Perhaps, then, instead of reporting an RPE, DA should be reinterpreted as reflecting the value of the animal’s current state, such as its position during reward approach (Hamid et al., 2016). Alternatively, perhaps DA signals different quantities in different tasks, e.g., value in operant tasks, in which the animal must act to receive reward, and RPE in classical conditioning tasks, in which the animal need not act to receive reward.

To distinguish among these possibilities, we recently devised an experimental paradigm that dissociates the value and RPE interpretations of DA (Kim et al., 2020). We began with the insight that, in the experiments considered above, RPEs can be approximated as the derivative of value under the TD learning framework (Gershman (2014); see Methods). This implies that, to effectively arbitrate between the value and RPE interpretations, one only need devise experiments where value and its derivative are expected to behave very differently. Indeed, by training mice on a virtual reality environment and manipulating various properties of the task—namely, the speed of scene movement and the presence of forward teleportations and temporary pauses—we could make precise predictions about how value should change vs. how its derivative (RPE) should change. We found that the changes in DA behaviors were consistent with the RPE hypothesis and not with the value interpretation. The virtual reality task further allowed us to dissociate spatial navigation from locomotion (running), as one view of ramps had been that they are specific to operant tasks, and that DA conveys qualitatively different information in operant vs. classical conditioning tasks. However, we found that mice continued to display ramping DA signals during the task even without locomotion (i.e., when the mice did not run for reward). We confirmed these key results at the levels of somatic spiking of DA neurons, axonal calcium signals, and DA concentrations at neuronal terminals in striatum. Taken together, these findings strongly support the RPE hypothesis of DA.

The body of experimental studies outlined above produces a number of unanswered questions regarding the function of DA: First, why would an error signal persist once an association is well-learned? Second, why would it ramp over the duration of the trial? Third, why would this ramp occur in some tasks but not others? Does value (and thus RPE) take different functional forms in different tasks, and if so, what determines which forms result in a ramp and which do not? Here we address these questions from normative principles.

We begin this work by examining the influence of sensory feedback in guiding value estimation. Because of irreducible temporal uncertainty, animals not receiving sensory feedback (and therefore relying only on internal timekeeping mechanisms) will have corrupted value estimates regardless of how well a task is learned. In this case, value functions will be ‘blurred’ in proportion to the uncertainty at each point. Sensory feedback, however, reduces this blurring as each new timepoint is approached. Beginning with the normative principle that animals seek to best learn the value of each state, we show that unbiased learning, in the presence of feedback, requires RPEs that ramp. These ramps scale with the informativeness of the feedback (i.e., the reduction in uncertainty), and at the extreme, absence of feedback leads to flat RPEs. Thus we show that differences in a task’s feedback profile explain the puzzling collection of DA behaviors described above. To experimentally verify our hypothesis, we trained mice on a virtual navigation task in which the brightness of the virtual track was varied. As predicted by our framework, when the scene was darkened over the course of the trial (putatively decreasing the sensory feedback), DA exhibited a ‘bump,’ or a ramp-up followed by a ramp-down. Furthermore, the magnitude of signals during the ramp-up phase was globally greater than that of the corresponding ramp in conditions when the scene brightness remained high, as predicted by the theory.

We will begin the next section with a review of the TD learning algorithm, then examine the effect of state uncertainty on value learning. We will then show how, by reducing state uncertainty without biasing learning, sensory feedback causes the RPE to reproduce the experimentally observed behaviors of DA. Finally, we will specifically control the sensory feedback by manipulating the brightness of the track in a virtual navigation task, thereby uncovering DA bumps.

## Results

### Temporal Difference Learning

In TD learning, an agent transitions through a sequence of states according to a Markov process (Sutton, 1988). The value associated with each state is defined as the expected discounted future return:

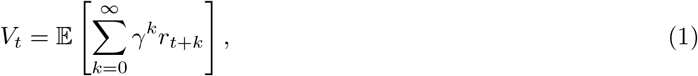

where *t* denotes time and indexes states, *r_t_* denotes the reward delivered at time *t*, and *γ* ∈ (0, 1) is a discount factor. In the experiments we will examine, a single reward is presented at the end of each trial. For these cases, Equation (1) can be written simply as:

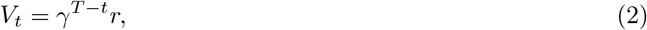

for all *t* ∈ [0, *T*], where *r* is the magnitude of reward delivered at time *T*. In words, value increases exponentially as reward time *T* is approached, peaking at a value of *r* at *T* (Figure 1B,D). Additionally, note that exponential functions are convex; the convex shape of the value function will be important in subsequent sections (see Kim et al. (2020) for an experimental test of this property).

**Figure 1:**
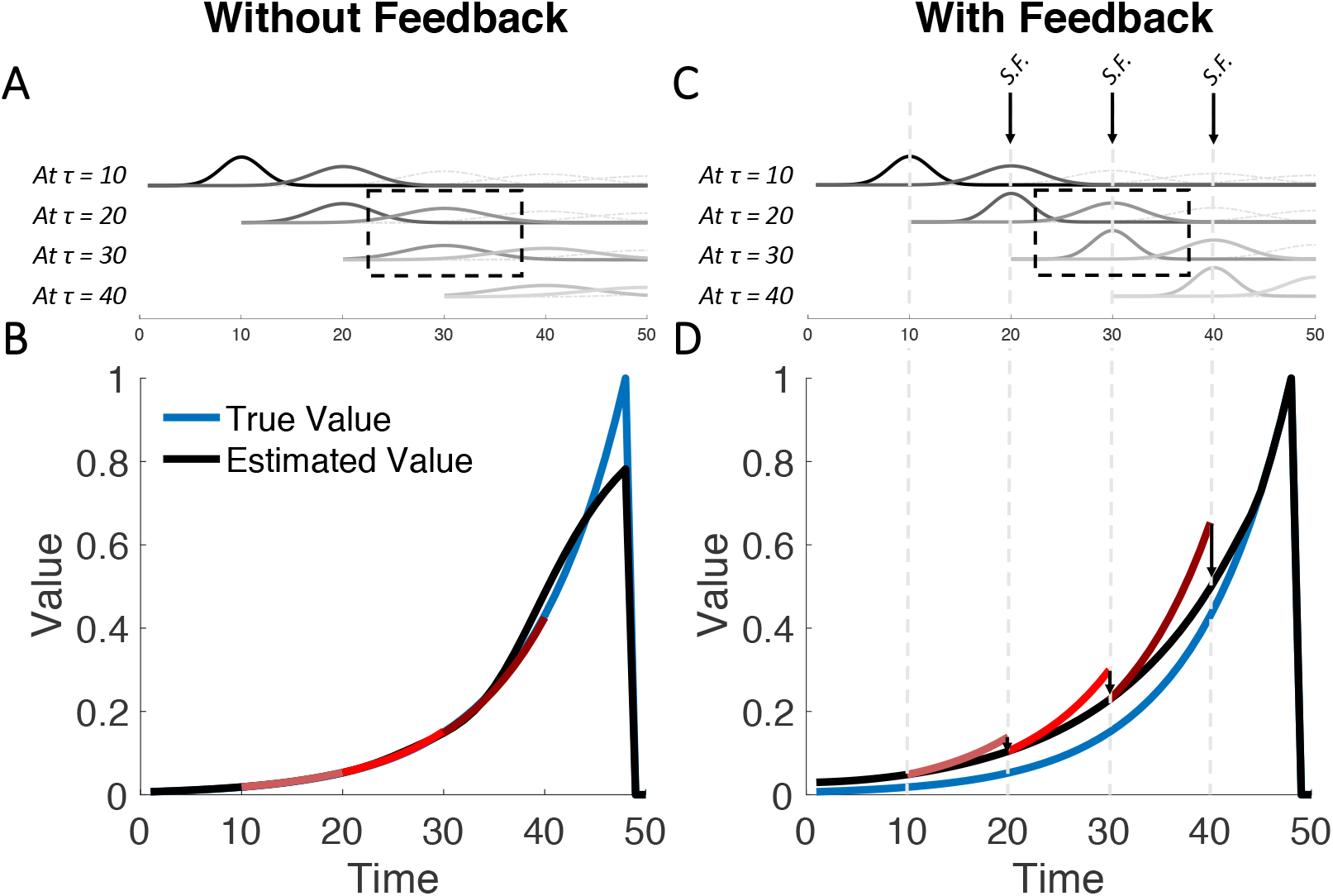
Sensory Feedback Biases Value Learning. (A) Illustration of state uncertainty in the absence of sensory feedback. Each row includes the uncertainty kernels at the current state and the next state (solid curves). Lighter gray curves represent uncertainty kernels for later states. Thus, similarly colored kernels on different rows represent uncertainty kernels for the same state, but evaluated at different timepoints (e.g., dashed box). In the absence of feedback, state uncertainty for a single state does not acutely change across time (compare with C). (B) Without feedback, value is unbiased on average. Red curves represent the predicted increase in value between the current state and the next state (10 and 20 for light red; 20 and 30 for red; 30 and 40 for brick red). After learning, this roughly equals an increase by *γ*^−1^ on average. (C) Sensory feedback reduces state uncertainty. Three instances of feedback are shown for illustration (*S.F.*; arrows). Note here that the kernels used to estimate value at the same state have different widths depending on whether they were evaluated before or after feedback. This results in different value estimates being used to compute the RPE at the current state and at the next state (Equations (8) and (9)). (D) As a result of sensory feedback, value at each state will be estimated based on an inflated version of value at the next state. Hence, after learning (when RPE is zero on average), estimated value will be systematically larger than true value. Red curves represent the predicted increase in value between the current state and the next state. After learning, this roughly equals an increase by *γ*^−1^ on average. See Methods for simulation details.

How does the agent learn this value function? Under the Markov property, the value at any time *t*, defined in Equation (1), can be rewritten as a sum of the reward received at *t* and the discounted value at the next time step:

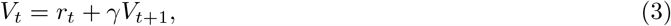

which is referred to as the Bellman equation (Bellman, 1957). In words, value at time *t* is the sum of rewards received at *t* and the promise of future rewards. To learn *V_t_*, the agent approximates it with 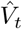, which is updated in the event of a mismatch between the estimated value and the reward actually received. By analogy with Equation (3), this mismatch (the RPE) can be written as:

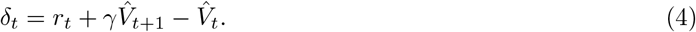

When *δ_t_* is zero, Equation (3) has been well-approximated. However, when *δ_t_* is positive or negative, 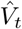 must be increased or decreased, respectively:

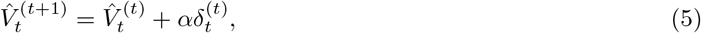

where *α* ∈ (0, 1) denotes the learning rate, and the superscript denotes the learning step. Learning will progress until *δ_t_* = 0 on average. After this point, 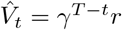 on average, which is precisely the true value. (See the Methods for a more general description of TD learning and its neural implementation.)

Having described TD learning in the simplified case where the agent has a perfect internal clock and thus no state uncertainty, let us now examine how state uncertainty affects learning, and how this uncertainty is reduced with sensory feedback.

### Value Learning Under State Uncertainty

Because animals do not have perfect internal clocks, they do not have complete access to the true time *t* (Gibbon, 1977; Church and Meck, 2003; Staddon, 1965). Instead, *t* is a latent state corrupted by timing noise, often modeled as follows:

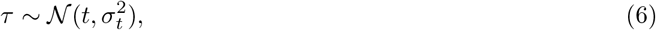

where *τ* is subjective time, drawn from a distribution centered on objective time *t*, with some standard deviation *σ_t_*. We take this distribution to be Gaussian for simplicity (an assumption we relax in the Methods). Thus the subjective estimate of value 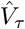 is an average over the estimated values 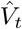 of each state *t*:

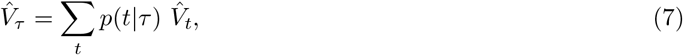

where *p*(*t*|*τ*) denotes the probability that *t* is the true state given the subjective measurement *τ*, and thus represents state uncertainty. We refer to this quantity as the uncertainty kernel (Figure 1A,C). Intuitively, 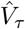 is the result of blurring 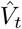 proportionally to the uncertainty kernel (Methods).

After learning (i.e., when the RPE is zero on average), the estimated value at every state will be roughly the estimated value at the next state, discounted by *γ*, on average (black curve in Figure 1B). A key requirement for this unbiased learning can be discovered by writing the RPE equations for two successive states:

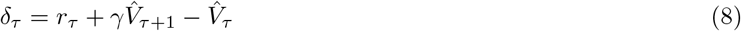

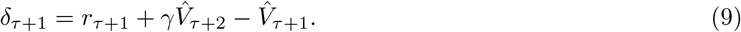

Notice here that 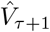 is represented in both equations. In other words, 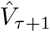 must be computed at two separate timepoints: at *τ* (where it represents the value of the next state) and at *τ* + 1 (where it represents the value of the new, current state). The TD equations, in their standard form, require that 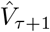 remain the same regardless of when it is computed, to achieve unbiased value-learning. Said differently, for value to be well-learned, a requirement is that 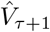 not acutely change during the interval after computing *δ_τ_* and before computing *δ_τ_*_+1_. This requirement extends to changes in the uncertainty kernels: By Equation (7), if the kernel *p*(*t*|*τ* + 1) were to be acutely updated due to information available at *τ* + 1 but not at *τ*, then 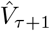 will acutely change as well. This means 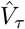 that will be discounted based on 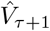 before feedback (i.e., as estimated at *τ*; red curves in Figure 1D) rather than 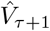 after feedback (i.e., as estimated at *τ* + 1; black curve). In the next section, we will examine this effect more precisely, and we will show that any such acute change (here, due to sensory feedback) will cause an unbiased agent to produce ramping RPEs.

### Value Learning in the Presence of Sensory Feedback

How is value learning affected by sensory feedback? As each time *τ* is approached, state uncertainty is reduced due to sensory feedback (arrows in Figure 1C). This is because at timepoints preceding *τ*, the estimate of what the value *will be* at *τ* is corrupted by both temporal noise and the lower-resolution stimuli associated with *τ*. Approaching *τ* in the presence of sensory feedback reduces this corruption. This, however, means that 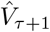 will be estimated differently while computing *δ_τ_* and *δ_τ_*+1 (Equations (8) and (9); compare widths of similarly shaded kernels beneath each arrow in Figure 1C)—a violation of the requirement mentioned above, which in turn results in biased value learning.

To examine the nature of this bias, we note that averaging over a convex value function results in over-estimation of value. Intuitively, convex functions are steeper on the right (larger values) and shallower on the left (smaller values), so averaging results in a bias toward larger values. Furthermore, wider kernels result in greater overestimation (Methods). Thus upon entering each new state, the reduction of uncertainty via sensory feedback will acutely mitigate this overestimation, resulting in different estimates 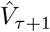 being used for *δ_τ_* and *δ_τ_*_+1_. Left uncorrected, the value estimate will be systematically biased, and in particular, value will be overestimated at every point (Figure 2A; Methods). An intuitive way to see this is as follows: The objective of the TD algorithm (in this simplified task setting) is for the value at each state *τ* to be *γ* times smaller than the value at *τ* + 1 by the time the RPE converges to zero (Equation (2)). If an animal systematically overestimates value at the next state, then it will overestimate value at the current state as well (even if sensory feedback subsequently diminishes the next state’s overestimation). Thus the ‘wrong’ value function is learned (Figure 2A,B).

**Figure 2:**
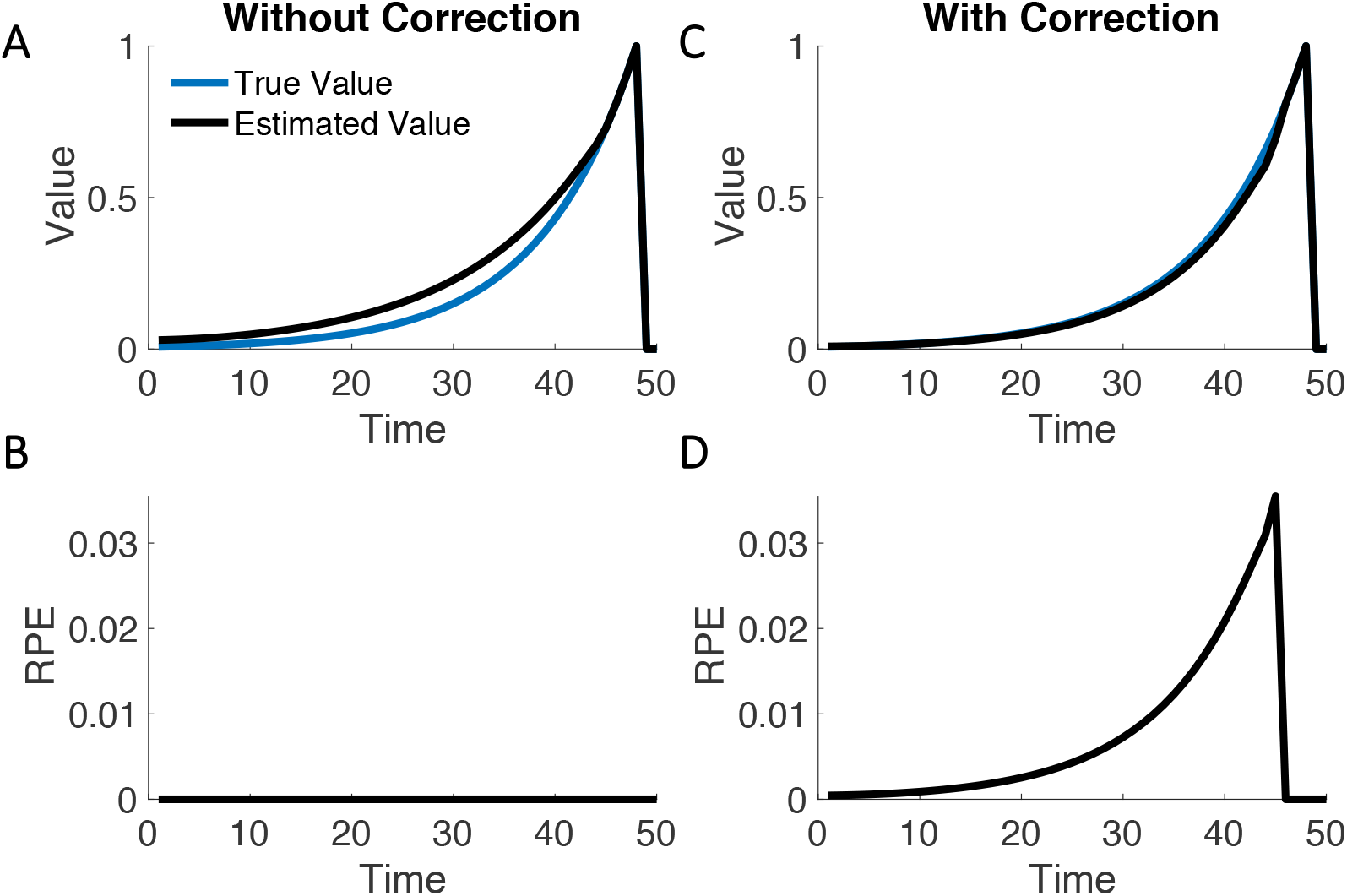
Unbiased Learning in the Presence of Feedback Leads to RPE Ramps. (A) In a hypothetical task with sensory feedback but in which correction does not occur, value at each state is learned according to an overestimated version of value at the next state. Thus, a biased (suboptimal) value function is learned (see Figure 1D). (B) After learning, the RPE converges to zero. (C) With a correction term, the correct value function is learned instead. (D) The cost of forcing an unbiased learning of value is a persistent RPE. Intuitively, value at the current state is not influenced by the overestimated version of value at the next state (compare with A,B). By Equation (13), this results in RPEs that ramp. See Methods for simulation details.

To overcome this bias, an optimal agent must correct the just-computed RPE as sensory feedback becomes available. In the Methods, we show that this correction can simply be written as:

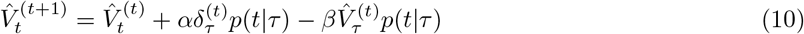

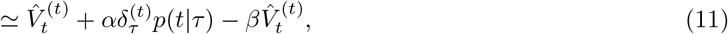

where the approximate equality holds for sufficient reductions in state uncertainty due to feedback, and

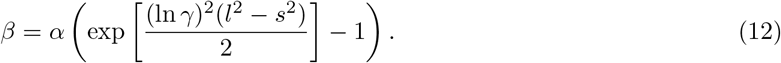

Here, the uncertainty kernel of 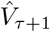 has some standard deviation *l* at *τ* and a smaller standard deviation *s* at *τ* + 1. In words, as the animal gains an improved estimate of 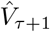, it corrects the previously computed *δ_τ_* with a feedback term to ensure unbiased learning of value (Figure 2C). Notice here that the correction term is a function of the reduction in variance (*l*^2^ − *s*^2^) due to sensory feedback. In the absence of feedback, the reduction in variance is zero (the uncertainty kernel for *τ* + 1 cannot be reduced during the transition from *τ* to *τ* + 1), which means *β* = 0.

How does this correction affect the RPE? By Equation (10), the RPE will converge to:

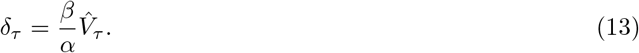

Therefore, with sensory feedback, the RPE ramps and tracks 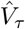 in shape (Figure 2D). In the absence of feedback, *β* = 0; thus, there is no ramp. Note here that the RPE is not a function of the learning rate *α*, as *β* itself is directly proportional to *α* (Equation (12)).

In summary, when feedback is provided with new states, value learning becomes miscalibrated, as each value point will be learned according to an overestimated version of the next (Figure 2A). With a subsequent correction of this bias, the agent will continue to overestimate the RPEs at each point (RPEs will ramp; Figure 2D), in exchange for learning the correct value function (Figure 2C).

### Relationship with Experimental Data

In classical conditioning tasks without sensory feedback, DA ramping is not observed (Schultz et al., 1997; Kobayashi and Schultz, 2008; Stuber et al., 2008; Flagel et al., 2011; Cohen et al., 2012; Hart et al., 2014; Eshel et al., 2015; Menegas et al., 2015, 2017; Babayan et al., 2018) (Figure 3A). On the other hand, in goal-directed navigation tasks, characterized by sensory feedback in the form of salient visual cues as well as locomotive cues (e.g., joint movement), DA ramping is present (Howe et al., 2013) (Figure 3C). DA ramping is also present in classical conditioning tasks that do not involve locomotion but that include either spatial or non-spatial feedback (Kim et al., 2020), as well as in two-armed bandit tasks (Hamid et al., 2016), in the timing of movement initiation (Hamilos et al., 2020), and when executing self-initiated action sequences (Wassum et al., 2012; Collins et al., 2016).

**Figure 3:**
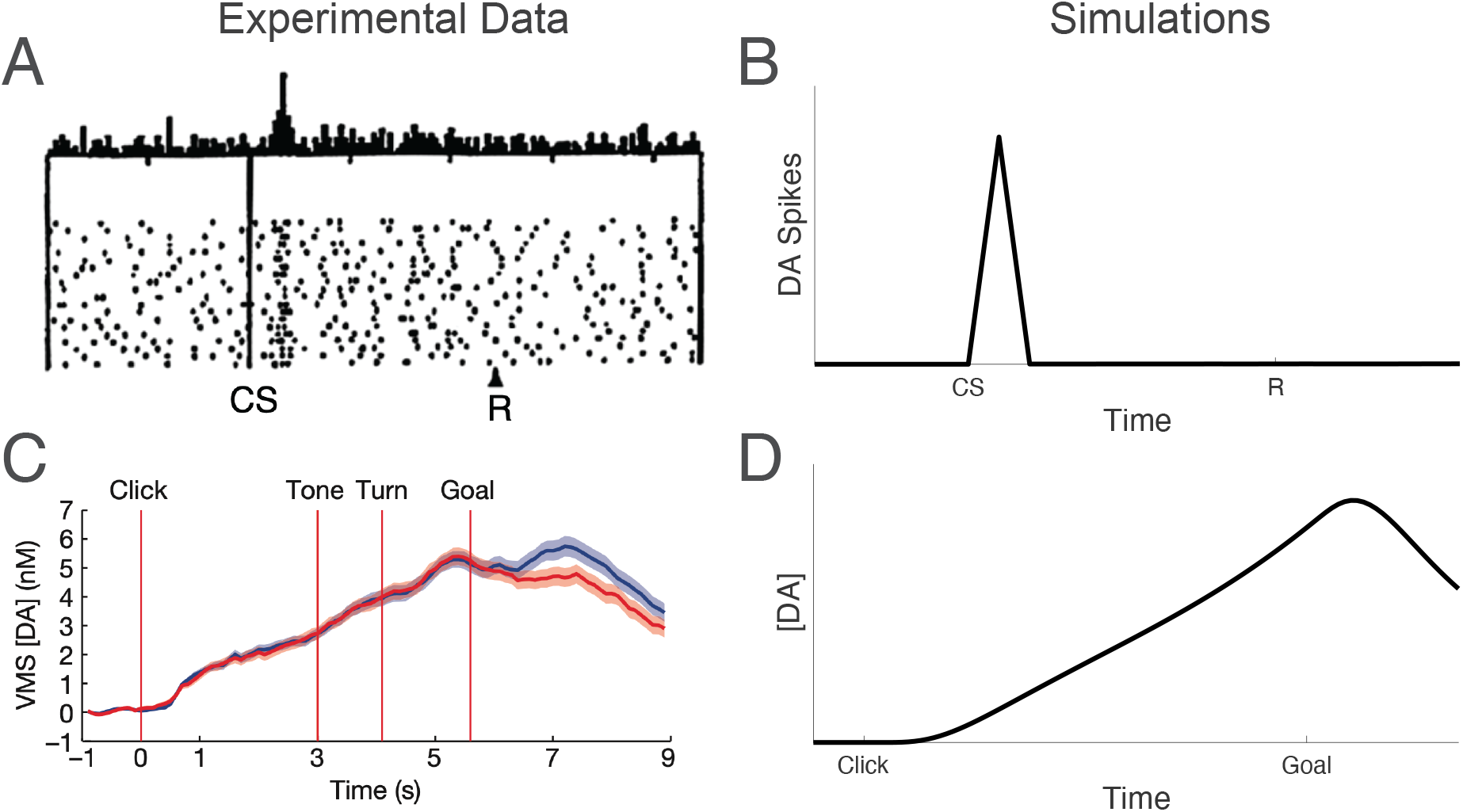
Differences in Feedback Result in Different RPE Behaviors. (A) Schultz et al. (1997) have found that after learning, phasic DA responses to a predicted reward (R) diminish, and instead begin to appear at the earliest reward-predicting cue (conditioned stimulus; CS). Figure from Schultz et al. (1997). (B) Our derivations recapitulate this result. In the absence of sensory feedback, RPEs converge to zero. Note here the absence of an RPE at reward time in the experimental data. This is predicted by the model because the CS-R duration is very small (under 1.5 seconds) in the experimental paradigm, so temporal uncertainty is also small. Longer durations are predicted to result in an irreducible RPE response, as has been experimentally observed (Kobayashi and Schultz, 2008), a point we return to in the Discussion. (C) Howe et al. (2013) have found that the DA signal ramps during a well-learned navigation task over the course of a single trial. Figure from Howe et al. (2013). (D) Our derivations recapitulate this result. In the presence of sensory feedback, RPEs track the shape of the estimated value function. See Methods for simulation details.

As described in the previous section, sensory feedback—due to external cues or to the animal’s own movement— can reconcile both types of DA behaviors with the RPE hypothesis: In the absence of feedback, there is no reduction in state uncertainty upon entering each new state (*β* = 0), and therefore no ramps (Equation (13); Figure 3B). On the other hand, when state uncertainty is reduced as each state is entered, ramps will occur (Figure 3D). Intuitively, information received after an RPE has already been computed (and hence, after a DA response has already occurred) biases the learning of value. To offset this bias, the RPE converges to be non-zero at the equilibrium state (when value is well-learned). Furthermore, because of the convexity of the value function, this non-zero RPE must increase as the reward is approached.

In a direct test of the competing views of DA, we recently devised a series of experiments to disentangle the value and RPE interpretations (Figure 4, top panels; Kim et al., 2020). We trained mice on a virtual reality paradigm, in which the animals experience virtual spatial navigation toward a reward. Visual stimuli on the (virtual) walls on either side of the path afforded the animals information about their location at any given moment. We then introduced a number of experimental manipulations—changing the speed of virtual motion, introducing a forward ‘teleportation’ at various start and end points along the path, and pausing the navigation for 5 seconds before resuming virtual motion. We showed that the value interpretation of DA made starkly different predictions from the RPE hypothesis, and then demonstrated that DA behavior was consistent with RPEs and not values.

**Figure 4:**
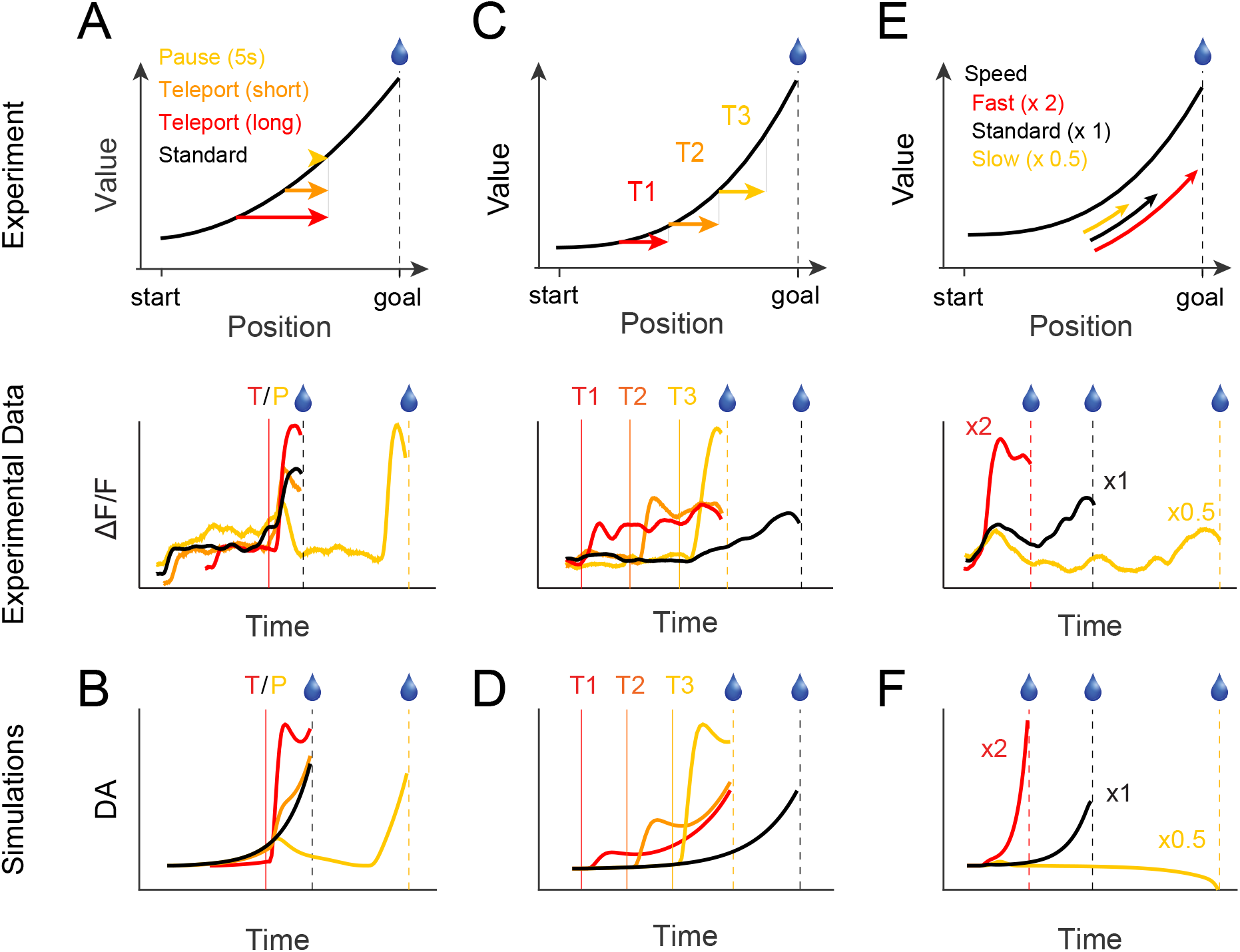
RPE Behaviors Match DA Responses Under Various Task Manipulations. We trained head-fixed mice on a visual virtual reality task, in which they virtually navigated a scene with a reward at the end (Kim et al., 2020). We then manipulated various aspects of the task. (A) When the mice were teleported from different locations to the same end point, a large DA response resulted, and scaled with the size of the teleport. When the navigation was paused for 5 seconds, the DA response dropped to baseline, with a large response occurring upon resuming navigation. (B) Our derivations recapitulate this result. With an instantaneous jump toward the reward, the RPE is very large, and increases with larger jumps. During a pause, the RPE drops to zero, but rapidly increases when navigation resumes. (C) When the mice were teleported from different locations but with the same magnitude, large DA responses resulted, and increased in size closer to the reward. (D) Our derivations recapitulate this result. Because of the convexity of the value function, an instantaneous teleportation of fixed magnitude will result in a larger RPE when it occurs closer to the reward. (E) When the scene was navigated through more quickly, the ramp was steeper. (F) Our derivations recapitulate this result. Faster navigation results in denser visual feedback per timepoint, i.e., the uncertainty kernels, defined by visual landmarks, become tighter with respect to true time. By Equations (12) and (13), this results in a greater reduction in uncertainty, and thus a steeper ramp. Panels (A,C,E) from Kim et al. (2020). See Methods for simulation details.

To show this difference, we noted that RPEs can be approximated as the derivative of value (Equation (4), where *r_t_* = 0 leading up to reward time, and *γ* is close to 1; note that this view ignores any contribution of state uncertainty). We then assumed that value is ‘sufficiently convex’ (Methods), in order to produce a derivative that increases monotonically. The task, then, was to simply examine the expected effect of each experimental manipulation on value vs. its derivative.

This view is limited in a number of ways. Perhaps most importantly, the presented model—that RPEs are the approximate derivative of value—fails to capture the recursive effect of RPEs on value: Not only does a value estimate generate an RPE, but the RPE also modifies the value estimate. If RPEs ramp, then they are always positive. But how, then, can the agent settle on a single value estimate, if the RPE is always causing the estimate to increase? A second limitation of this model is that it had to *assume* a sufficiently convex value function, in order to achieve a monotonically increasing derivative (and hence a ramping RPE), leaving open the question of where this convexity originates from. Finally, this view cannot accommodate experiments where ramps are not observed. Instead, the model would seemingly predict ramping in all tasks, even though, as amply discussed above, this is not the case (e.g., Schultz et al., 1997; Kobayashi and Schultz, 2008). In Figure 4, we show that our uncertainty-based model, which is not subject to these limitations, predicts the entire range of experimental results in Kim et al. (2020).

### Manipulation of Sensory Feedback and DA Bumps

We have shown that our framework captures an array of DA behaviors. However, the manipulations considered above do not isolate sensory feedback as the key contributor to ramping. We therefore sought to develop an experimental paradigm that can distinguish our uncertainty-based model from the conventional models.

By describing a relationship between sensory feedback and DA ramps, our model predicts that a wide variety of DA responses can be elicited under the appropriate uncertainty profiles. In particular, our model makes an interesting prediction about a third type of behavior that to our knowledge has not been previously observed: If state uncertainty rapidly increases over the course of a trial, then rather than a ramp, DA responses should exhibit a bump (Figure 5D). To see this intuitively, we can examine the RPE behaviors early and late in a trial in which the visual scene is gradually darkened, putatively decreasing the sensory feedback over the course of the trial. Initially, when the brightness is still high, the RPE should behave as in the constant-brightness condition (i.e., ramps). As the scene darkens, wider uncertainty kernels ‘blur’ the convex value function more. Thus the early ramp in the darkening condition will be higher than that of the constant condition. However, later in the trial, as the animal approaches the reward, wider uncertainty kernels serve to flatten the estimated value function (near the maximum value, averaging over a larger window decreases the value estimate). Thus the RPE will begin to decrease. Taken together, this results in an RPE bump that increases early on and decreases later. Furthermore, because of the lack of feedback near the reward time, the flatter estimated value function will result in a larger reward response than in the constant condition.

**Figure 5:**
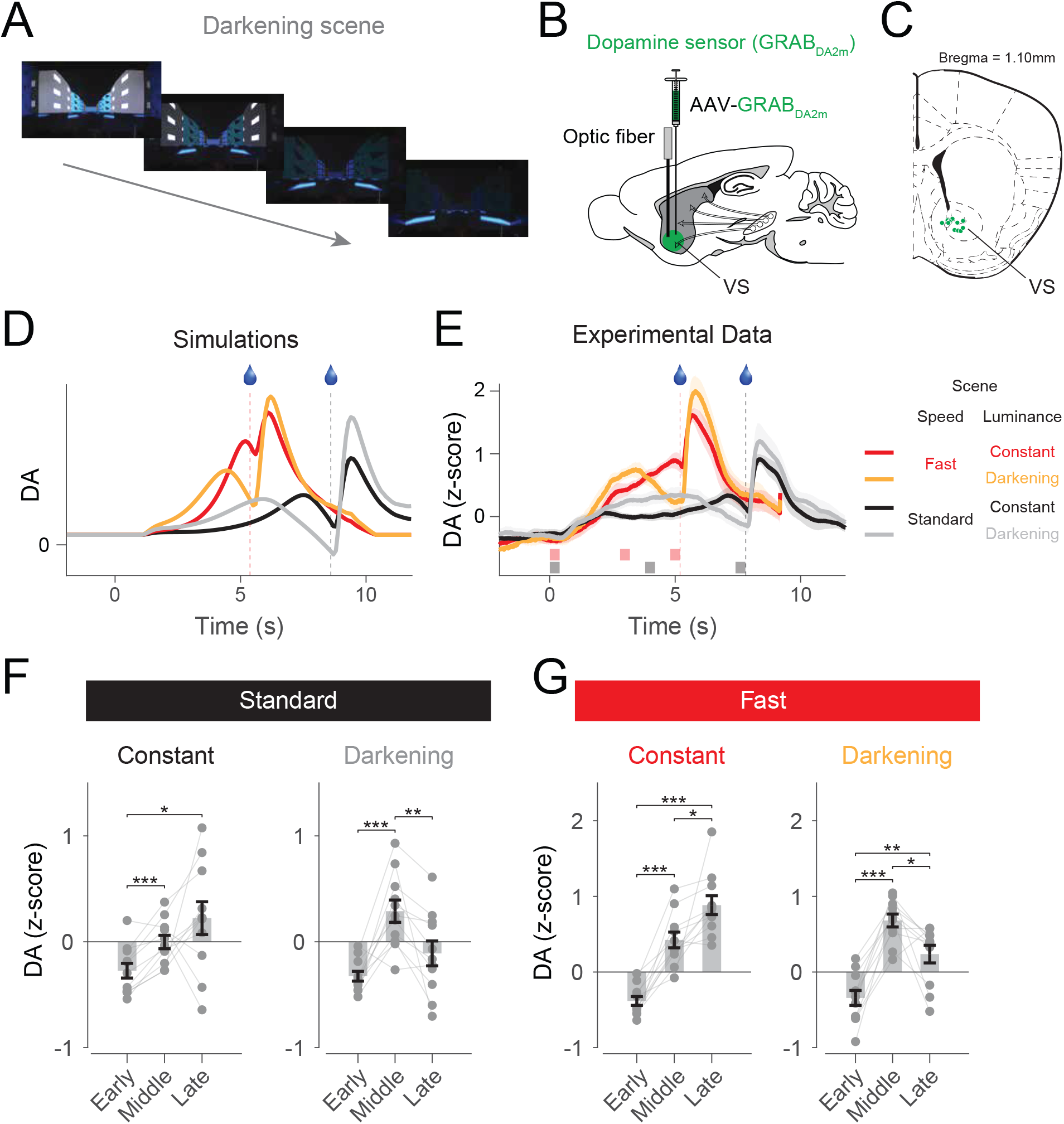
The State Uncertainty Model Predicts DA Responses in the Darkening Experiments. (A) Images of the visual scene captured at four different locations. The floor patterns were intact to prevent animals from inferring that the trial was aborted. (B) Experimental design for fiber fluorometry. Adeno-associated virus (AAV) expressing a DA sensor (GRAB_DA2m_) was injected into the ventral striatum (VS). DA signals were monitored through an optical fiber implanted into the VS. (C) Recording locations. A coronal section of the brain at Bregma, 1.10 mm. (D) Model predictions. Note three properties of the DA response in the darkening condition: the DA bump, the greater initial ramp compared to the constant condition, and the stronger reward response compared to the constant condition. Black, constant condition with standard speed; gray, darkening condition with standard speed; red, constant condition with fast speed (x1.7); yellow, darkening condition with fast speed. (E) DA responses. Shaded areas at the bottom depict time windows for the three epochs used in (F,G). (F) Average DA responses in the standard conditions. Three dots connected with lines represent individual animals (*n* = 11 mice). (G) Average DA responses in the fast conditions (*n* = 11 mice). Shadings and error bars represent standard errors of the mean. ^*^*p* < 0.05, ^**^*p* < 0.01, ^***^*p* < 0.001, *t*-test.

In order to test these predictions explicitly, we dynamically modulated the reliability of sensory evidence by changing the brightness of the visual scene over the course of a single trial (‘darkening’ condition; Figure 5; Figure S3; Supplemental Movie 1). The darkening condition (25% of trials) was randomly interleaved with the constant-brightness condition (75% of trials). We independently interleaved the standard-speed and fast conditions (on 25% of trials, the scene moved 1.7 times faster than the standard-speed condition). Including a small portion of fast conditions appeared to help animals pay attention to the task. We monitored DA activity in the ventral striatum using fiber fluorometry (Figure 5B,C). Note that animals showed anticipatory licking in the darkening conditions (Figure S3B), suggesting that the animals did not think the trials were aborted.

As predicted, our manipulations of scene brightness—putatively manipulations of the sensory feedback— caused a DA bump, a signal that increases early on and decreases later (Figure 5E, gray and yellow curves). When the scene moved at standard speed, DA activity modestly ramped up in the constant condition (Figure 5F, left), whereas DA activity displayed a bump in the darkening condition (Figure 5F, right). The average responses in the middle epoch were significantly greater than those of either the start or end epoch (*p* < 0.01, *t*-test, *n* = 11 mice). Ramping in the constant condition became more evident when the scene moved fast (Figure 5G, left). Nevertheless, we still observed a bump in the middle when the visual scene was darkened (Figure 5G, right). Furthermore, because of the lack of feedback near the reward time, our model predicts that the flatter estimated value function will result in a larger (phasic) response to the reward, compared to the constant condition, for both the standard and fast conditions, as indeed observed (Figure S3C, left and right, respectively; *p* < 0.01, *t*-test, *n* = 11 mice).

## Discussion

While a large body of work has established phasic DA as an error signal (Schultz et al., 1997; Niv and Schoenbaum, 2008; Glimcher, 2011; Steinberg et al., 2013; Eshel et al., 2015), more recent work has questioned this view (Wassum et al., 2012; Howe et al., 2013; Hamid et al., 2016; Collins et al., 2016). Indeed, in light of persistent DA ramps occurring in certain tasks even after extensive learning, some authors have proposed that DA may instead communicate value itself in these tasks (Hamid et al., 2016). However, the determinants of DA ramps have remained unclear: Ramps are observed during goal-directed navigation, in which animals must run to receive reward (operant tasks; Howe et al., 2013), but can also be elicited in virtual reality tasks in which animals do not need to run for reward (classical conditioning tasks; Kim et al., 2020). Within classical conditioning, DA ramps can occur in the presence of navigational or non-navigational stimuli indicating time to reward (Kim et al., 2020). Within operant tasks, ramps can be observed in the period preceding the action (Totah et al., 2013) as well as during the action itself (Howe et al., 2013). These ramps are furthermore not specific to experimental techniques and measurements, and can be observed in cell body activities, axonal calcium signals, and in the DA concentrations (Kim et al., 2020).

We have shown in this work that, under the RPE hypothesis of DA, sensory feedback may control the different observed DA behaviors: In the presence of persistent sensory feedback, RPEs track the estimated value in shape (ramps), but they remain flat in the absence of feedback (no ramps). Thus DA ramps and phasic responses follow from common computational principles and may be generated by common neurobiological mechanisms. Moreover, a curious lemma of this result is that a measured DA signal whose shape tracks with estimated value need not be evidence against the RPE hypothesis of DA, contrary to some claims (Hamid et al., 2016; Berke, 2018): Indeed, in the presence of persistent sensory feedback, *δ_τ_* and 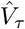 have the same shape. Thus, our derivation is conceptually compatible with the value interpretation of DA under certain circumstances, but importantly, this derivation captures the experimental findings in other circumstances in which the value interpretation fails (see below for further discussion).

Our model implies that a variety of peculiar DA responses can be attained under the appropriate sensory feedback profiles. In particular, knowing that value increases monotonically over the course of a trial, our results imply that a rapidly decreasing sensory feedback profile will result in a previously unobserved DA bump. By testing animals on conditions in which the visual scenes gradually darkened over the course of a single trial, we found exactly this result: a DA response that ramps up early on and ramps down later.

Our work takes inspiration from previous studies that examined the role of state uncertainty in DA responses (Kobayashi and Schultz, 2008; Fiorillo et al., 2008; de Lafuente and Romo, 2011; Starkweather et al., 2017; Lak et al., 2017). For instance, temporal uncertainty increases with longer durations (Staddon, 1965; Gibbon, 1977; Church and Meck, 2003). This means that in a classical conditioning task, DA bursts at reward time will not be completely diminished, and will be larger for longer durations, as Kobayashi and Schultz (2008) and Fiorillo et al. (2008) have observed. Similarly, Starkweather et al. (2017) have found that in tasks with uncertainty both in *whether* reward will be delivered as well as *when* it is delivered, DA exhibits a prolonged dip (i.e., a negative ramp) leading up to reward delivery. Here, value initially increases as expected reward time is approached, but then begins to slowly decrease as the probability of reward delivery during the present trial becomes less and less likely, resulting in persistently negative prediction errors (see also Starkweather et al., 2018; Babayan et al., 2018). As the authors of these studies note, both results are fully predicted by the RPE hypothesis of DA. Hence, state uncertainty, due to noise either in the internal circuitry or in the external environment, is reflected in the DA signal.

### Alternative Hypotheses

One might argue that state uncertainty is not necessary to explain the results in the darkening experiments. To address this issue, we considered the possibilities that the DA responses can be explained either by the value interpretation of DA or by an RPE hypothesis that does not account for state uncertainty (Supplemental Text 1). Briefly, the non-monotonic behavior of the DA response is incompatible with the value interpretation of DA, as darkening the visual scene should not decrease the value. Indeed, the animals’ lick rates continued to increase in both the constant and darkening conditions (Figure S3). Second, the DA patterns are incompatible with the conventional, uncertainty-independent RPE view. To show this, we recovered the value functions from the putative RPE signals, and found that the value in the darkening condition would have to be globally greater than that in the constant condition. However, under the uncertainty-free RPE hypothesis, value in the darkening condition should either be the same as in the constant condition (value estimates unaffected by brightness) or smaller (if an inability to see the reward at the end of the trial leads to an assumed reward probability that is less than 1). We expand on these points in Supplemental Text 1.

Finally, we note that our results are based on the assumption that animals maintain the same value function across experimental conditions. Said differently, we have assumed here that animals learn the value function in the constant condition and subsequently apply this previously learned value function to probe trials in which the scene is gradually darkened. It is possible, however, that animals learn a separate value function for the darkening conditions. Because RPEs in our model increase with larger values and decrease with lower feedback, it remains possible that such an alternative model will still capture the observed effects (Supplemental Text 2).

While we have derived RPE ramping from normative principles, it is important to note that a complete correction is not necessary to produce ramping. Furthermore, biases in value learning may also produce ramping. For instance, one earlier proposal by Gershman (2014) was that value may take a fixed convex shape in spatial navigation tasks; the mismatch between this shape and the exponential shape in Equation (2) produces a ramp (see Methods for a general derivation of the conditions for a ramp). Morita and Kato (2014), on the other hand, posited that value updating involves a decay term, which is qualitatively similar to that in Equation (10), and thus RPE ramping (see also implementations in Mikhael and Bogacz, 2016; Cinotti et al., 2019). Ramping can similarly be explained by assuming temporal or spatial bias that decreases with approach to the reward, by modulating the temporal discount term during task execution, or by other mechanisms (Supplemental Text 4). In each of these proposals, ramps emerge as a ‘bug’ in the implementation, rather than as an optimal strategy for unbiased learning. These proposals furthermore do not explain the different DA patterns that emerge under different paradigms. Finally, it should be noted that we have not assumed any modality- or task-driven differences in learning (any differences in the shape of the RPE follow solely from the sensory feedback profile), although in principle, different value functions may certainly be learned in different types of tasks (e.g., Supplemental Text 2).

Alternative accounts of DA ramping that deviate more significantly from our framework have also been proposed. In particular, Lloyd and Dayan (2015) have provided three compelling theoretical accounts of ramping. In the first account, the authors show that within an actor-critic framework, uncertainty in the communicated information between actor and critic regarding the timing of action execution may result in a monotonically increasing RPE leading up to the action. In the second account, ramping modulates gain control for value accumulation within a drift-diffusion model (e.g., by modulating neuronal excitability; Nicola et al., 2000). Under this framework, fluctuations in tonic and phasic DA produce average ramping. The third account extends the average reward rate model of tonic DA proposed by Niv et al. (2007). In this extended view, ramping constitutes a ‘quasi-tonic’ signal that reflects discounted vigor. The authors show that the discounted average reward rate follows (1 − *γ*)*V*, and hence takes the shape of the value function in TD learning models. Ramps may also result from *perceived* control, i.e., they may only occur if the animal *thinks* it can control the outcome of the task. While the Kim et al. (2020) virtual reality experiments strongly argue against this possibility, as the head-fixed animals who did not display running behavior during the task still exhibited ramps, it remains possible that these animals adopted some other, unmeasured superstitious behavior, thus resulting in perceived control. Finally, and relatedly, Howe et al. (2013) have proposed that ramps may be necessary for sustained motivation in the operant tasks considered. Indeed, the notion that DA may serve multiple functions beyond the communication of RPEs is well-motivated and deeply ingrained (Schultz, 2007b, 2010; Berridge, 2007; Frank et al., 2007; Gardner et al., 2018). Our work does not necessarily invalidate these alternative interpretations, but rather shows how a single RPE interpretation can embrace a range of apparently inconsistent phenomena.

### Lingering Questions

A number of questions arise from our analysis. First, is there any evidence to support the benefits of learning the ‘true’ value function as written in Equation (2) (Figure 2C) over the biased version of value (Figure 2A)? We note here that under the normative account, the agent seeks to learn *some* value function that maximizes its well-being, whose exact shape has been the subject of much interest (e.g., Rachlin and Green, 1972; Ainslie, 1975; Tobin and Logue, 1994; Rachlin, 2000). Our key result is that this function—regardless of its exact shape—will not be learned well if feedback is delivered during learning, unless correction ensues. Beyond learning a suboptimal value function, the agent will furthermore be biased *across* options, as two equally rewarding options will generate different value functions if one was learned with feedback and the other was not (see Methods for a similar case in which this bias is costly). Note also that, while we have chosen the exponential shape in Equation (2) after the conventional TD models, our ramping results extend to any convex value function.

Second, due to the presumed exponential shape, the ramping behaviors resulting from our analysis may also at times look exponential, rather than linear. We nonetheless have chosen to remain close to conventional TD models and purely exponential value functions for ease of comparison with the existing theoretical literature. Perhaps equally important, the relationship between RPE and its neural correlate need only be monotonic and not necessarily equal. In other words, a measured linear signal does not necessarily imply a linear RPE, and a convex neural signal need not communicate convex information. It remains an open question how best to bring abstract TD models into alignment with biophysically realistic assumptions about the signal-generating process.

## Methods

### Temporal Difference Learning and Its Neural Correlates

Under TD learning, each state is determined by task-relevant contextual cues, referred to as features, that predict future rewards. For instance, a state might be determined by a subjective estimate of time or perceived distance from a reward. We model the agent as approximating *V_t_* by taking a linear combination of the features (Schultz et al., 1997; Ludvig et al., 2008, 2012):

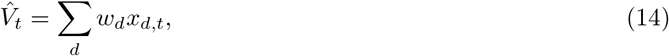

where 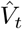 denotes the estimated value at time *t*, and *x_d,t_* denotes the *d^th^* feature at *t*. The learned relevance of each feature *x_d_* is reflected in its weight *w_d_*, and the weights are updated in the event of a mismatch between the estimated value and the rewards actually received. The update occurs in proportion to each weight’s contribution to the value estimate at *t*:

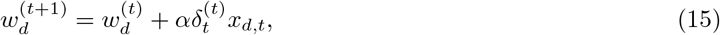

where *α* ∈ (0, 1) denotes the learning rate, and the superscript denotes the learning step. In words, when a feature *x_d_* does not contribute to the value estimate at *t* (*x_d,t_* = 0), its weight is not updated. On the other hand, weights corresponding to features that do contribute to 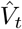 will be updated in proportion to their activations at that time. This update rule is referred to as gradient ascent (*x_d,t_* is equal to the gradient 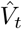 of with respect to the weight *w_d_*), and it implements a form of credit assignment, in which the features most activated at *t* undergo the greatest modification to their weights.

In this formulation, the basal ganglia implements the TD algorithm termwise: Cortical inputs to striatum encode the features *x_d,t_*, corticostriatal synaptic strengths encode the weights *w_d_* (Houk et al., 1995; Montague et al., 1996), phasic activity of midbrain DA neurons encodes the error signal *δ_t_* (Schultz et al., 1997; Niv and Schoenbaum, 2008; Glimcher, 2011; Steinberg et al., 2013; Eshel et al., 2015), and the output nuclei of the basal ganglia (substantia nigra pars reticulata and internal globus pallidus) encode estimated value 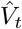 (Ratcliff and Frank, 2012).

We have implicitly assumed in the Results a maximally flexible feature set, the complete serial compound representation (Moore et al., 1989; Sutton and Barto, 1990; Montague et al., 1996; Schultz et al., 1997), in which every time step following trial onset is represented as a separate feature. In other words, the feature *x_d,t_* is 1 when *t* = *d* and 0 otherwise. In this case, value at each timepoint is updated independently of the other timepoints, and each has its own weight. It follows that 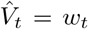, and we can write Equation (15) directly in terms of 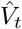, as in Equation (5).

### Value Learning Under State Uncertainty

The animal has access to subjective time *τ*, from which it forms a belief state *p*(*t*|*τ*), or, in Bayesian terms, a posterior distribution over true time. For simplicity, we have taken this distribution to be Gaussian, and we assume weak priors so that temporal estimates, though noisy, are accurate. In this case, the subjective time estimate 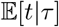 is and is equal to the posterior mean. Note here that we are only concerned with capturing the noisy property of internal clocks. While a large literature has sought to establish the exact relationship between internal (‘psychological’) time and true time with varying degrees of success (e.g., linear vs. logarithmic relationship; Allan, 2002; Wearden, 2002; Wearden and Jones, 2007; Jozefowiez et al., 2018; Ren et al., 2020), our work is invariant to this exact relationship, and only depends on animals’ ability to reproduce time veridically on average, with some noise (Gibbon, 1977; Church and Meck, 2003; Staddon, 1965).

Given the subjective time *τ*, the RPE is then:

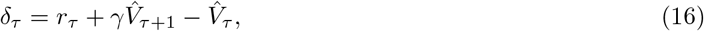

and this error signal is used to update the value estimates at each point *t* in proportion to its posterior probability *p*(*t*|*τ*):

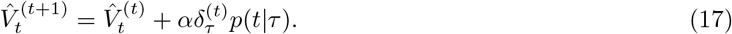

Said differently, the effect of state uncertainty is that when the error signal *δ_τ_* is computed, it updates the value estimate at a number of timepoints, in proportion to the uncertainty kernel.

Note here that, in the absence of uncertainty, our task structure obeys the Markov property: state transitions and rewards are independent of the animal’s history given its current state. An appeal of using belief states is that the task remains Markovian, but in the posterior distributions rather than in the signals, and the TD algorithm can be applied directly to our learning problem, as in Equations (16) and (17). This problem is a type of partially observable Markov decision process (Gershman and Uchida, 2019).

### Acute Changes in State Uncertainty Result in Biased Value Learning

Averaging over a convex value function results in overestimation of value. For an exponential value function, we can derive this result analytically in the continuous time domain:

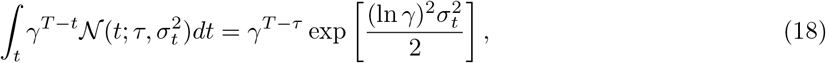

where *σ_t_* is the standard deviation of the uncertainty kernel at *t*, and the second term on the right-hand side is greater than one. Intuitively, because the function is steeper on the right side and shallower on the left side, the average will be overestimated. Importantly, however, the estimate will be a multiple of the true value, with a scaling factor that depends on the width of the kernel (second term on right-hand side of Equation (18); note also that while we have assumed a Gaussian distribution, our qualitative results hold for any distribution that results in overestimation of value). Thus, with sensory feedback that modifies the width of the kernel upon transitioning from one state (*τ*) to the next (*τ* + 1), there will be a mismatch in the value estimate when computing each RPE. More precisely, the learning rules are:

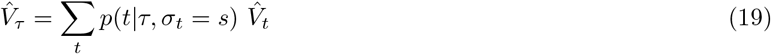

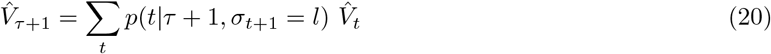

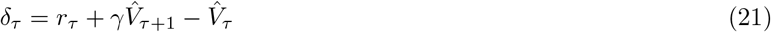

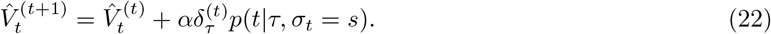

Notice that 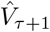 takes different values depending on the state: When computing *δ_τ_*,

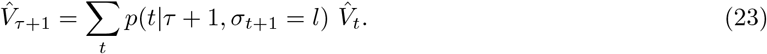

On the other hand, when computing *δ_τ_*_+1_,

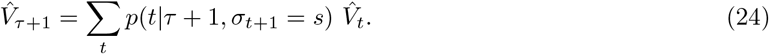

How does this mismatch affect the learned value estimate? If averaging with kernels of different standard deviations can be written as multiples of true value, then they can be written as multiples of each other. The RPE is then

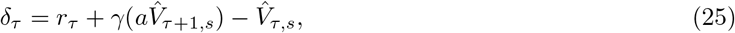

where we use the comma notation in the subscripts to denote that the two value estimates are evaluated with the same kernel width *s*, and *a* is a constant. By analogy with Equations (2) and (4), estimated value converges to 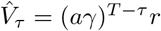. Here, *a* > 1, so value is systematically overestimated. By the learning rules in Equations (19) to (22), this is because *δ_τ_* is inflated by

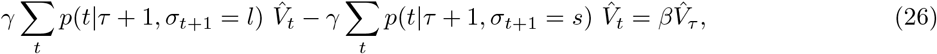

where *β* is defined in Equation (12).

An optimal agent will use the available sensory feedback to overcome this biased learning. Because averaging with a kernel of width *l* is simply a multiple of that with width *s*, it follows that a simple subtraction can achieve this correction (Equations (10) and (11)). Hence, sensory feedback can improve value learning with a correction term. It should be noted that with a complete correction to *s* as derived above, the bias is fully extinguished. For corrections to intermediate widths between *s* and *l*, the bias will be partially corrected but not eliminated. In both cases, because *β* > 0, ramps will occur.

In extension of the first Methods section, we can posit an implementation of uncertainty kernels in which sensory information is relayed from cortical areas (Houk et al., 1995; Montague et al., 1996) and the uncertainty due to Weber’s law is based in fronto-striatal circuitry (Matell et al., 2005).

### RPEs Are Approximately the Derivative of Value

Consider the formula for RPEs in Equation (4). In tasks where a single reward is delivered at *T*, *r_t_* = 0 for all *t < T* (no rewards delivered before *T*). Because *γ* ≃ 1, the RPE can be approximated as

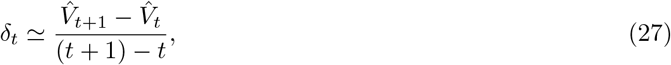

which is the slope of the estimated value. To examine the relationship between value and RPEs more precisely, we can extend our analysis to the continuous domain:

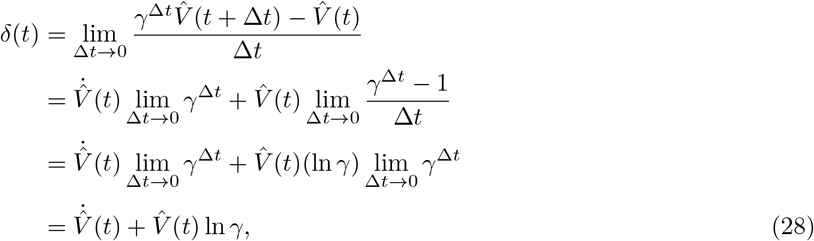

where 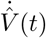 is the time derivative of 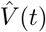 and the third equality follows from L’Hôpital’s Rule. Here, ln *γ* has units of inverse time. Because ln *γ* ≃ 0, RPE is approximately the derivative of value.

### Sensory Feedback in Continuous Time

In the complete absence of sensory feedback, *σ_t_* is not constant, but rather increases linearly with time, a phenomenon referred to as *scalar variability*, a manifestation of Weber’s law in the domain of timing (Gibbon, 1977; Church and Meck, 2003; Staddon, 1965). In this case, we can write the standard deviation as *σ_t_* = *wt*, where *w* is the Weber fraction, which is constant over the duration of the trial.

Set *l* = *w*(*τ* + Δ*τ*) and *s* = *wτ*. Following the steps in the previous section,

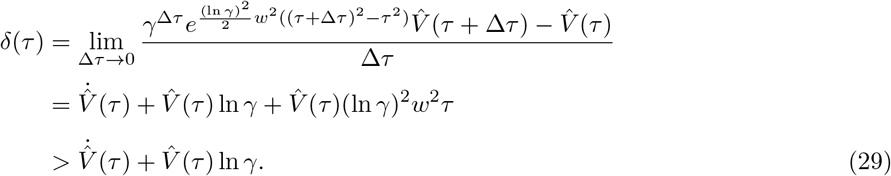

Hence, as derived for the discrete case, RPEs are inflated, and value is systematically overestimated.

### RPE Ramps Result From Sufficiently Convex Value Functions

By Equation (28), the condition for ramping is 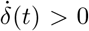 i.e., the estimated shape of the value function at any given point, before feedback, must obey

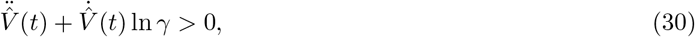

where 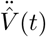 is the second derivative of 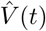 with respect to time. For an intuition of this relation, note that when *γ* ≃ 1, the inequality can be approximated as 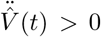 which denotes any convex function. The exact inequality, however, has a tighter requirement on 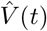: Since 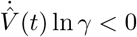 for all *t*, ramping will only be observed if the contribution from 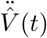 (i.e., the convexity) outweighs the quantity 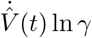 (the scaled slope). For example, the function in Equation (2) does not satisfy the strict inequality even though it is convex, and therefore with this choice of 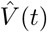, the RPE does not ramp. In other words, to produce an RPE ramp 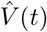, has to be ‘sufficiently’ convex.

### Biased Value Estimates and Reward Forfeiture

Let us illustrate here how a biased value function can lead to suboptimal choices. Imagine a two-armed bandit task in which the animal chooses between two options, *A* and *B*, yielding rewards *rA* and *rB*, respectively, after a fixed delay *T*.

Assume *r_A_* = 1 is learned under conditions with rich sensory feedback, and *r_B_* = 1.5 is learned without feedback. Assume, also, that the animal learns according to the TD algorithm without a correction term. Using the simulation parameters for Figure 2A, with a delay of *T* = 20, it follows that the values at the time of choice are 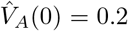 (Figure 2A, black curve at *t* = 28) and 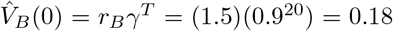 (Figure 2A, approximated as blue curve at *t* = 28, scaled by *r_B_*). After learning, the animal will be more likely to select *A*. (Furthermore, a greedy animal will asymptotically only select *A*.) With each selection of *A*, the animal forfeits an additional 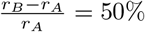 of reward potential.

### Simulation Details

#### Value Learning Under State Uncertainty (Figure 1)

For our TD learning model, we have chosen *γ* = 0.9, *α* = 0.1, *n* = 50 states, and *T* = 48. In the absence of feedback, uncertainty kernels are determined by the Weber fraction, set to *w* = 0.15 (Gallistel et al., 2004). In the presence of feedback, uncertainty kernels have a standard deviation of *l* = 3 before feedback and *s* = 0.1 after feedback. For the purposes of averaging with uncertainty kernels, value peaks at *T* and remains at its peak value after *T*, and the standard deviation at the last 4 states in the presence of feedback is fixed to 0.1. Intuitively, the animal expects reward to be delivered, and attributes any lack of reward delivery at *τ* = *T* to noise in its timing mechanism (uncertainty kernels have nonzero width) rather than to a reward omission. The learning rules were iterated 1000 times.

#### Value Learning in the Presence of Sensory Feedback (Figure 2)

For our TD learning model, we have chosen *γ* = 0.9, *α* = 0.1, *n* = 50 states, and *T* = 48. The learning rules were iterated 1000 times.

#### GCaMP Impulse Response Function

To model experiments involving Ca^2+^ signals, we used the GCaMP impulse response function obtained in Kim et al. (2020). This function was convolved with the computed RPEs to obtain simulated Ca^2+^ signals.

For convolutions over negative RPEs, it is important to account for the low baseline firing rates of DA neurons, i.e., that negative RPEs cannot elicit phasic responses that equal those elicited by positive RPEs of similar magnitude. Thus, following previous experimental (Bayer and Glimcher, 2005; Morris et al., 2004; Fiorillo et al., 2003) and theoretical (Daw et al., 2006, 2002; Niv et al., 2005) work, we account for an asymmetry between positive and negative RPEs in the DA signal. We do so by scaling the RPEs by the maximum change in spiking activity in either the positive or negative direction. After Kim et al. (2020), resting state spiking activity is approximately 5 spikes/second, the maximum spiking is 30 spikes/second, and the minimum spiking is 0 spikes/second. Thus one unit of positive RPE influences the DA response 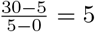 times as strongly as one unit of negative RPE.

#### Relationship with Experimental Data (Figures 3 and 4)

##### Figure 3

For our TD learning model, we have chosen *γ* = 0.98, *α* = 0.1, and Weber fraction *w* = 0.15. For the navigation task, kernels have standard deviation *l* = 3 before feedback and *s* = 0.1 after feedback. For and (D), we have set *n* = 10 and 70 states, respectively, between trial start and reward. The learning rules were iterated 2000 times.

##### Figure 4

The simulations of these experiments inherited the properties of the navigation task in Figure 3. For our TD learning model, we have chosen *γ* = 0.93, *α* = 0.1, *w* = 0.15, and *n* = 200 states. Kernels have standard deviation *l* = 1 before feedback and *s* = 0.5 after feedback for the teleport and pause manipulations, and *l* = 3 before feedback and *s* = 1 after feedback for the speed manipulation. The learning rules were iterated 2000 times.

#### Manipulation of Sensory Feedback and DA Bumps (Figure 5)

The TD model is identical to that in Figure 4. For the constant condition, the small kernel width is a constant, *s* = *a*. For the darkening condition, the width resembles that of the constant condition early on and resembles one without feedback later, (*s* − *a*)(*s* − *wt* − *b*) = *c*. The shape of this function is controlled by two parameters, *c* and *b*. The first determines how smoothly *s* transitions from resembling that of the constant condition to behaving according to Weber’s law, and the second determines when this occurs. The large uncertainty kernel width is *l* = *s* + *z*, where *z* is a constant in the constant condition, and *z* decreases smoothly to zero over the course of the trial in the darkening condition, which we model as 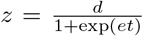. We set *a* = 8, *b* = 0.3, *c* = 3, *d* = 0.8, and *e* = 1.

### Subject Details

In addition to the fifteen GCaMP mice used in the previous study (Kim et al., 2020), eleven adult C57/BL6J wild-type male mice were used for the scene darkening experiments using the DA sensor. All mice were backcrossed for more than 5 generations with C57/BL6J mice. Animals were singly housed on a 12 hr dark/12 hr light cycle (dark from 07:00 to 19:00). All procedures were performed in accordance with the National Institutes of Health Guide for the Care and Use of Laboratory Animals and approved by the Harvard Animal Care and Use Committee.

### Surgery and Virus Injections

#### Surgery for Fiber Fluorometry of DA Sensor Signals

To prepare animals for recording, we performed a single surgery with three key components: (1) injection of a DA sensor into the ventral striatum, (2) head-plate installation, and (3) implantation of an optical fiber into the striatum (Babayan et al., 2018; Menegas et al., 2017). At the time of surgery, all mice were 2–4 months old. All surgeries were performed under aseptic conditions with animals anesthetized with isoflurane (1-2% at 0.5-1.0 L/min). Analgesia (ketoprofen for post-surgery treatment, 5 mg/kg, I.P.; buprenorphine for pre-operative treatment, 0.1 mg/kg, I.P.) was administered for 3 days following each surgery. We removed the skin above the surface of the brain and dried the skull using air. We injected 400 nL of AAV9-hSyn-DA2m (Vigene Biosciences) into the ventral striatum (bregma 1.0, lateral 1.1, depths 4.2 and 4.1 mm). Virus injection lasted several minutes, and then the injection pipette was slowly removed over the course of several minutes.

We then installed a head-plate for head-fixation by gluing a head-plate onto the top of the skull (C&B Metabond, Parkell). We used ring-shaped head-plates to ensure that the skull above the striatum would be accessible for fiber implants. Finally, during the same surgery, we also implanted optical fibers into the ventral striatum. To do this, we first slowly lowered optical fibers (200 *μ*m diameter, Doric Lenses) into the striatum using a fiber holder (SCH 1.25, Doric Lenses). The coordinates we used for targeting were bregma 1.0, lateral 1.1, depth 4.1 mm. Once fibers were lowered, we first attached them to the skull with UV-curing epoxy (Thorlabs, NOA81), and then a layer of black Ortho-Jet dental adhesive (Lang Dental, IL). After waiting for fifteen minutes for this glue to dry, we applied a small amount of rapid-curing epoxy (A00254, Devcon) to attach the fiber cannulas to the underlying glue and head-plate. After waiting for fifteen minutes for the epoxy to cure, the surgery was completed.

#### Surgery for Fiber Fluorometry of GCaMP Signals in the Ventral Striatum

To examine axonal calcium signals of dopaminergic neurons in the ventral striatum, we injected AAV-FLEX-GCaMP into the midbrain of DAT-Cre mice (Kim et al., 2020). Surgical procedures up to virus injection were the same as the DA sensor injections described above. We unilaterally injected 250 nL of AAV5-CAG-FLEX-GCaMP6m (1 x 10^12^ particles/mL, Penn Vector Core) into both the ventral tegmental area (VTA) and substantia nigra pars compacta (SNc) (500 nL total). To target the VTA, we made a small craniotomy and injected the virus at bregma 3.1, lateral 0.6, depths 4.4 and 4.1 mm. To target SNc, we injected the virus at bregma 3.3, lateral 1.6, depths 3.8 and 3.6 mm.

### Virtual Reality Setup

Virtual environments were displayed on three liquid crystal display (LCD) monitors with thin frames (Kim et al., 2020). VirMEn software (Aronov and Tank, 2014) was used to generate virtual objects and render visual images using perspective projection. Mice were head-restrained at the center of three monitors. Mice were placed on a cylindrical styrofoam treadmill (diameter 20.3 cm, width 10.4 cm). The rotational velocity of the treadmill was encoded using a rotary encoder. The output pulses of the encoder were converted into continuous voltage signals using a customized Arduino program running on a microprocessor (Teensy 3.2). Water reward was given through a water spout located in front of the animal’s mouth. Licking tongue movements were monitored using an infrared sensor (OPB819Z, TT Electronics). Voltage signals from the rotary encoder and the lick sensor were digitized into a PCI-based data-acquisition system (PCIe-6323, National Instruments) installed on the visual stimulation computer. Timing and amount of water were controlled through a micro-solenoid valve (LHDA 1221111H, The Lee Company) and switch (2N7000, On Semiconductor). Analog output TTL pulse was generated from the visual stimulation computer to deliver reward to the animals.

### Virtual Linear Track Experiments

Animals were trained in a virtual linear track (see Kim et al. (2020) for details). The maze was composed of a starting platform and a corridor with walls on both sides. We first trained animals on the standard approach-to-target task to learn the association between target location and reward. Once the animals learned the task, we ran a series of tasks with test trials to examine the nature of DA signals. In this paper, we simulated three main experiments in the previous study (Figure 4; Kim et al., 2020). We typically ran each task for two consecutive days (with a zero- or one-day break). Unless otherwise noted, unexpected reward (5 *μ*L) was given during the inter-trial interval on 3-6% of trials.

#### Scene Darkening Manipulation

We dynamically modulated the reliability of sensory evidence by changing the brightness of the visual scene (Supplemental Movie 1). The brightness of the visual scene at each time point was determined by multiplying the original RGB color values with a time-varying multiplier. The multiplier *k*(*t*) is a function of the animal’s position as defined below (Figure S3A).

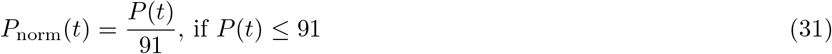

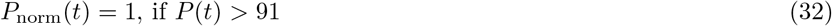

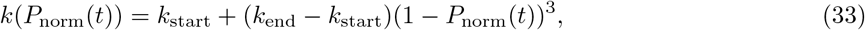

where *k*_start_ = 1.0, *k*_end_ = 0.05, and *P* (*t*) is animal’s position at time *t*. The brightness of the floor pattern was intact to provide the animals a clue that trials were not aborted. We randomly interleaved four experimental conditions. On 25% of trials, the visual scene was darkened as described above. Brightness was kept constant (*k*(*t*) = 1) for the rest of the trials. Independent of the brightness manipulation, the speed of visual scene progression was increased by 1.7 times on 25% of trials. Since the darkening depends on the position of the animal, for each darkening condition, the brightness of the scene at the reward location is identical between the standard and fast conditions.

### Fiber Fluorometry (Photometry)

Fluorescent signals from the brain were recorded using a custom-made fiber fluorometry (photometry) system as described in our previous studies (Kim et al., 2020; Menegas et al., 2017; Babayan et al., 2018). The blue light (473 nm) from a diode-pumped solid-state laser (DPSSL; 80–500 *μ*W; Opto Engine LLC, UT, USA) was attenuated through a neutral density filter (4.0 optical density, Thorlabs, NJ, USA) and coupled into an optical fiber patchcord (400 *μ*m, Doric Lenses) using a 0.65 NA microscope objective (Olympus). The patchcord connected to the implanted fiber was used to deliver excitation light to the brain and to collect the fluorescence emission signals from the brain. The fluorescent signal from the brain was spectrally separated from the excitation light using a dichroic mirror (T556lpxr, Chroma), passed through a bandpass filter (ET500/50, Chroma), focused onto a photodetector (FDS100, Thorlabs), and amplified using a current preamplifier (SR570, Stanford Research Systems). Acquisition from the red fluorophore (tdTomato) was simultaneously acquired (bandpass filter ET605/70 nm, Chroma) but was not used for further analyses. The voltage signal from the preamplifier was digitized through a data acquisition board (PCI-e6321, National Instruments) at 1 kHz and stored in a computer using a custom software written in LabVIEW (National Instruments).

### Histology

Mice were perfused with phosphate buffered saline (PBS) followed by 4% paraformaldehyde in PBS. The brains were cut in 100-*μ*m coronal sections using a vibratome (Leica). Brain sections were loaded on glass slides and stained with DAPI (Vectashield). The locations of fiber and tetrode tips were determined using the standard mouse brain atlas (Franklin and Paxinos, 2008).

### Quantification and Statistical Analysis

#### Statistical Analysis

We used a *t*-test to compare between conditions (Figure 5; Figure S1). Kolmogorov-Smirnov test was used to check the normality assumption.

#### Fluorometry (Photometry)

Power line noise in the raw voltage signals was removed by notch filter (MATLAB, Natick, MA, USA). A baseline of the voltage signal was defined by the lowest 10% of signals using a 2-min window. The baseline was subtracted from the raw signal, and the results were z-scored by a session-wide mean and standard deviation.

#### Licking and Locomotion

Lick timing was defined as deflection points (peaks) of the output signals above a threshold. To plot the time course of licks, instantaneous lick rate was computed by a moving average using a 200-ms window.

#### Session-Averaged Time Course

Licks, locomotion speed, and z-scored DA responses for individual trials were aligned by external events (e.g., trial start or teleport onset), and then smoothed using a moving average method. We did not smooth locomotion speed and fluorometry signals. The results were then averaged across trials for each experimental condition to generate a session-averaged time course.

#### Population-Averaged Time Course

For calcium recording experiments, we computed the mean of session-averaged time courses from the second session dataset (as the average of all session averages) along with the standard error (the total number of sessions being the sample size) for each experimental condition. Population-average time courses are used to summarize behavior and DA responses.

#### Quantification for the Darkening Experiments

We quantified the z-scored DA sensor responses in the darkening experiment using three time windows (Figure 5E, shaded areas at the bottom). For the standard conditions, we used [0 s 0.4 s] from the trial start, [3.8 s 4.2 s] from the trial start, and [-0.4 s 0 s] from the reward onset. For the fast conditions, we used [0 s 0.4 s] from the trial start, [2.8 s 3.2 s] from the trial start, and [-0.4 s 0 s] from the reward onset.

## Acknowledgments

The project described was supported by National Institutes of Health grants T32GM007753 and T32MH020017 (JGM), R01 MH110404 and MH095953 (NU), U19 NS113201-01 (SJG and NU), the Air Force Office of Scientific Research grant FA9550-20-1-0413 (SJG and NU), the Simons Collaboration on the Global Brain (NU), and a research fellowship from the Alfred P. Sloan Foundation (SJG). The content is solely the responsibility of the authors and does not necessarily represent the official views of the National Institutes of Health or the Simons Collaboration on the Global Brain. The funders had no role in study design, data collection and analysis, decision to publish, or preparation of the manuscript.

## Author Contributions

J.G.M. and S.J.G. developed the model. H.R.K. designed and conducted the experiments. H.R.K. and N.U. conceived that the structure of state uncertainty may influence the shape of estimated value functions and thus RPEs. J.G.M. analyzed and simulated the model. J.G.M., H.R.K., N.U., and S.J.G. contributed to the writing of the paper.

## Data and Code Availability

Data will be released upon publication. Source code for all simulations can be found at www.github.com/jgmikhael/ramping.

## Supplemental Information

### 1 Alternative Hypotheses and DA Bumps

We have argued in the main text that DA bumps can be captured by an uncertainty-driven view of RPEs but not by the value interpretation or the standard, uncertainty-free RPE hypothesis. To rule out the alternative hypotheses, we begin by deconvolving the GCaMP response, yielding a signal that we interpret as either pure value or uncertainty-free RPE.

The deconvolved signal is monotonic in the constant condition but non-monotonic in the darkening condition (Figure S1B). On the other hand, the licking data—putatively reflecting the animal’s estimate of value— increases monotonically in both conditions (Figure S3B, top panel). Taken together, these findings rule out the value interpretation of DA.

Next, we show that this signal is incompatible with an uncertainty-free RPE. To do so, we infer the value from the computed RPE (Figure S1C, using the derivation below). There is one free parameter, *γ*. We find that value is greater in the darkening condition than in the constant condition, even though under the uncertainty-free RPE hypothesis, it should either be the same (value estimate unaffected by brightness) or smaller (if an inability to see the reward location suggests a probability of receiving reward that is no longer equal to 1). Although *γ* is a free parameter, this result does not depend on *γ*, as 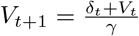, so *γ* simply amplifies or reduces existing differences, but does not reverse them.

To derive value from RPEs and *γ*, we use the relation:

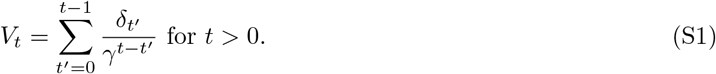

To show that Equation (S1) solves for *V_t_* using Equation (4) leading up to reward (i.e., when *r_t_* = 0), we use proof by induction. First, for *t* = 1,

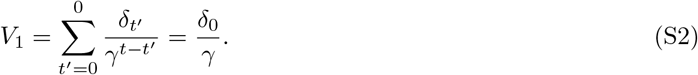

**Figure S1:**
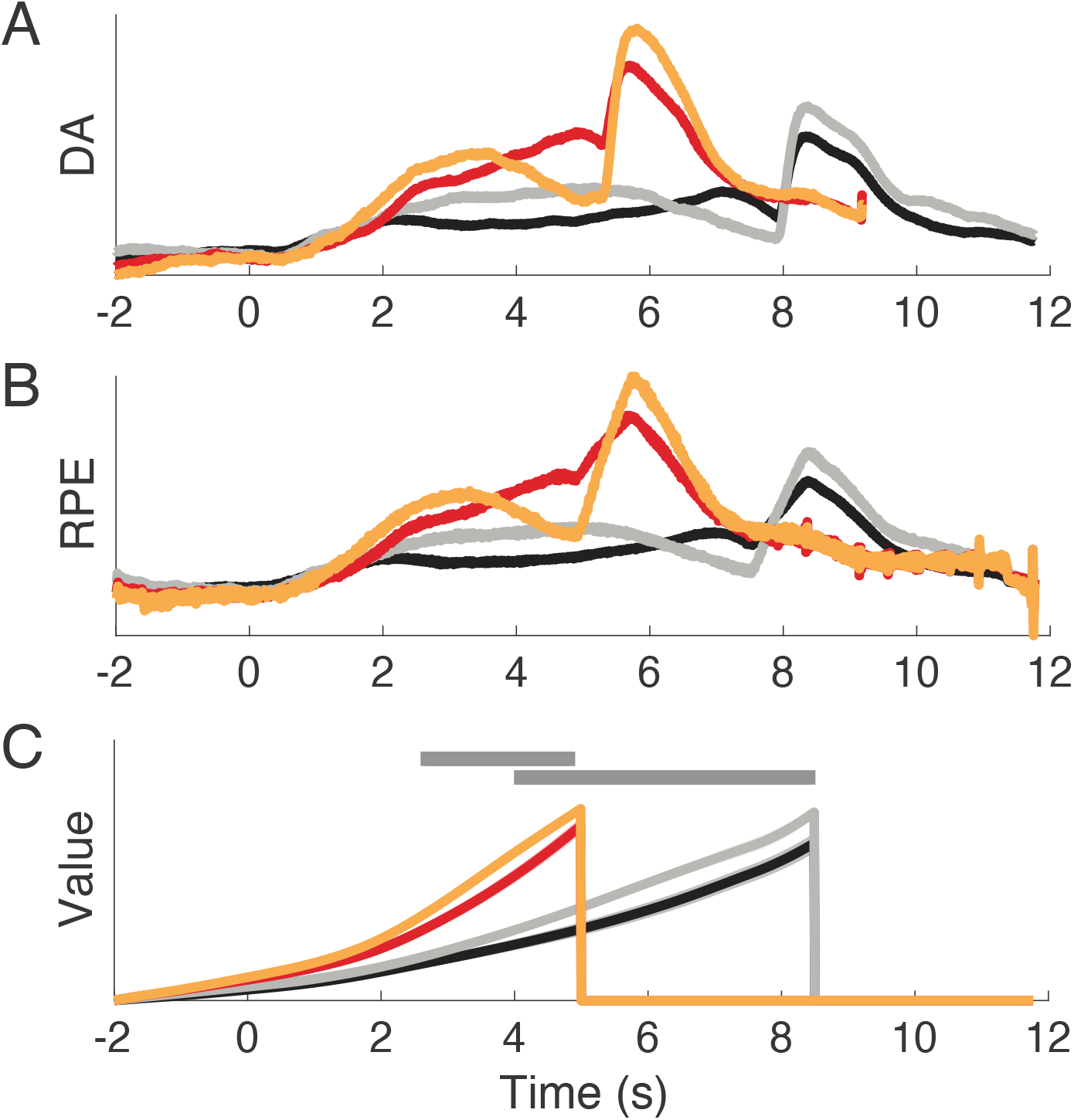
DA Bumps Are Incompatible With the Value or Uncertainty-Free RPE Views. (A) GCaMP responses. (B) Smoothed deconvolution of GCaMP responses, using an arbitrary smoothing window of approximately 0.2 seconds. (C) Under the assumption that the deconvolved signal represents RPE, we can derive the value. Value in the darkening condition is globally greater than value in the constant condition. For each animal, value is normalized in the fast and standard conditions separately. Gray horizontal bars represent statistically significant difference between the two conditions after the start of the scene movement (top: fast; bottom: standard). *p* < 0.05, *t*-test.

Thus Equation (S1) holds for *t* = 1. Now assume it holds for *t*; let us show it also holds for *t* + 1:

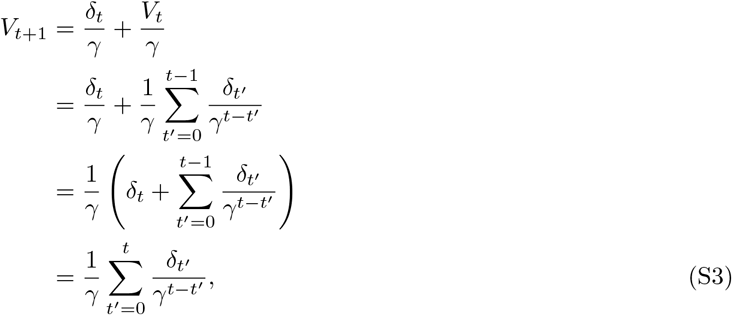

as required.

### 2 DA Bumps as a Consequence of Learning

In modeling the darkening manipulation, we have assumed that animals do not learn a separate value function for the probe trials in the darkening condition. We noted, however, that because of the opposite effects of the uncertainty profile and value on the RPE signal, bumps should still be observed when the manipulation occurs during learning (rather than only during performance). We show this analytically here.

Consider a manipulation in which the scene is gradually darkened, transitioning from perfect brightness to complete darkness over the course of a single trial. Using the terminology in the main text, the reduction in standard deviation (*l* − *s*) decreases monotonically over the course of the trial (less sensory feedback), eventually reaching zero. But value increases monotonically over the trial, starting at zero. By Equation (13), the RPE reflects a product of 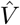 and *β*, which itself depends on (*l*^2^ −*s*^2^) = (*l* −*s*)(*l* +*s*). This means that the RPE should be zero at the beginning of the task and the end, but be positive in the middle. Because both *V* and *β* are continuous and differentiable, so is their product. Thus we predict that the RPE will gradually increase, reach some maximum, and subsequently decrease back to zero within a single trial (Figure S2).

**Figure S2:**
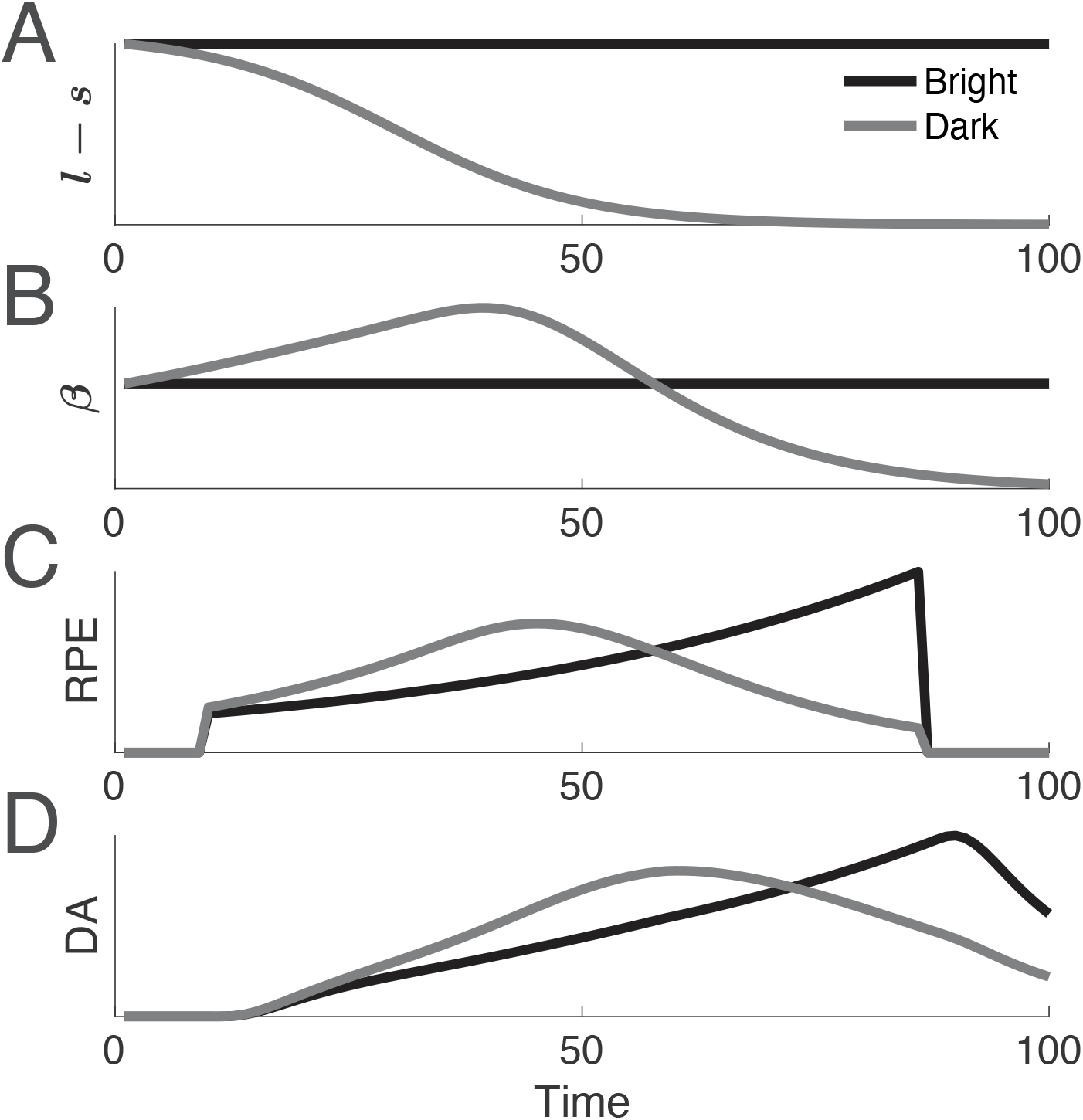
Illustration of the Learning Model for the Darkening Condition. (A) Temporal profiles of the difference in the widths of the two uncertainty kernels. In the constant condition, correction due to sensory feedback is constant throughout the trial. In the darkening condition, the correction decreases with time. (B) Temporal profiles of *β*, which is proportional to (*l*^2^ - *s*^2^) = (*l − s*)(*l* + *s*). (C) Temporal profiles of the RPE, which is proportional to the product of 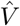 and *β*. (D) RPEs convolved with GCaMP impulse response function.

### 3 Experimental Details and Behaviors in the Darkening Condition

**Figure S3:**
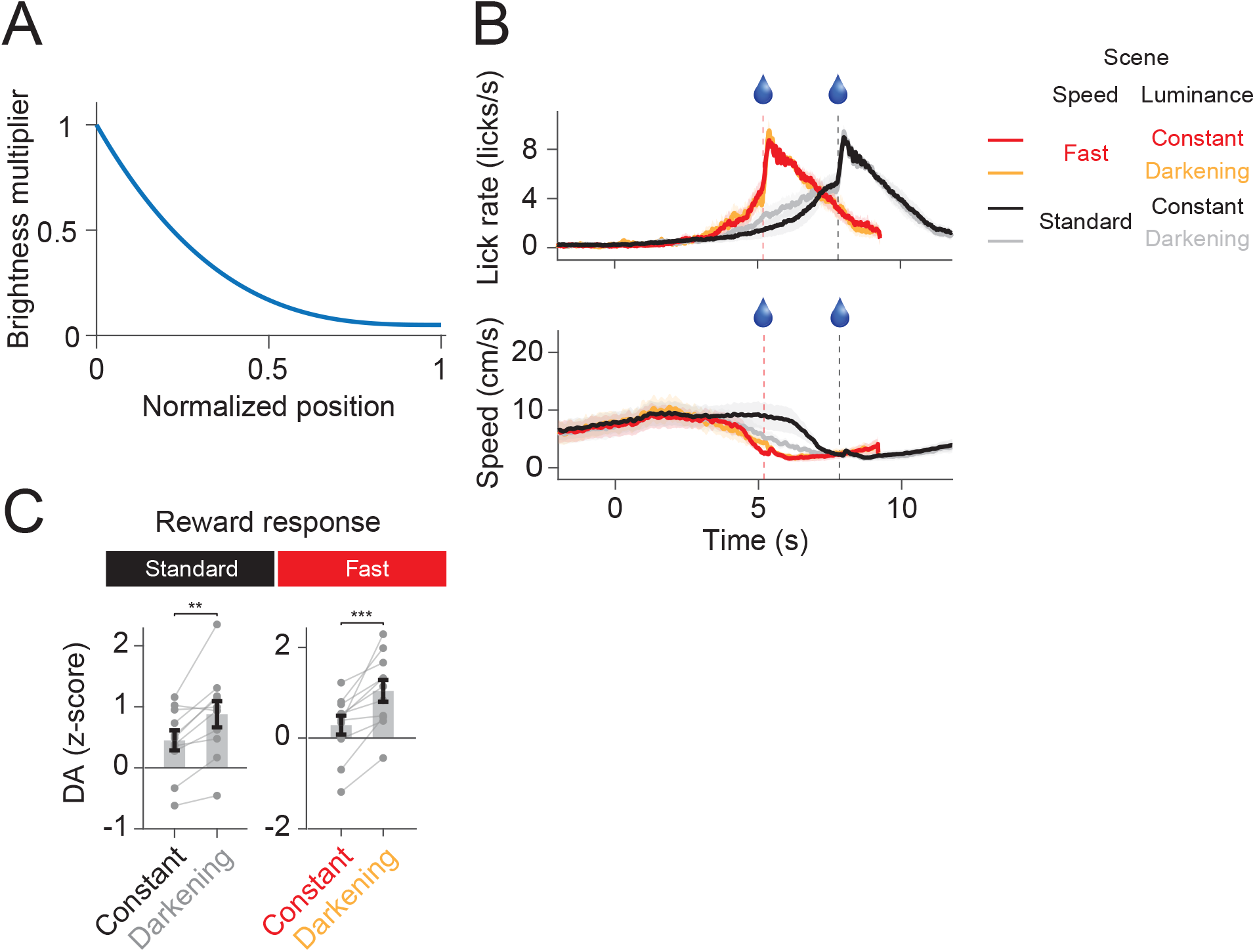
Experimental Details and Behaviors in the Darkening Condition. (A) Brightness multiplier as a function of the animal’s normalized position. (B) Average lick rate (top) and locomotion speed (bottom) (*n* = 11 mice). (C) Reward response magnitudes (average response 0-2 s from reward onset). DA activity at the time of reward delivery was subtracted from reward responses. Shadings and error bars represent standard errors of the mean. ^**^*p* < 0.01, ^***^*p* < 0.001, *t*-test.

### 4 Alternative Causes of Ramping

In the main text, we argued that ramping follows from normative principles. In this section, we illustrate that various types of biases (‘bugs’ in the implementation) may also lead to RPE ramps.

#### Ramping Due to Bias in State Estimation

Assume the animal persistently overestimates the amount of time or distance remaining to reach its reward (or, equivalently, that it underestimates the time elapsed or the distance traversed so far), and that this overestimation decreases as the animal approaches the reward. For instance, since the receptive fields of place cells decrease as the animal approaches reward (O’Keefe and Burgess, 1996), the contribution of place cells immediately behind the approaching animal in its estimate of value may outweigh that from the place cells in front of it. It will simplify our analysis to set *T* = 0 without loss of generality, and allow time to progress from the negative domain (*t* < 0) toward *T* = 0. In the continuous domain and for the simple case of linear overestimation, we can write this as

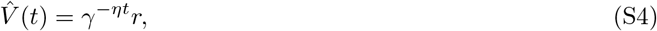

where *η* > 1 is our overestimation factor. Therefore, by Equation (28),

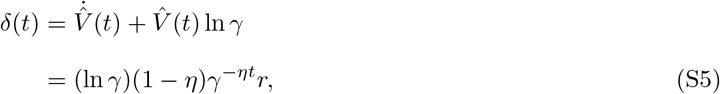

which is monotonically increasing. Hence, the RPE should ramp. Equivalently, in the discrete domain,

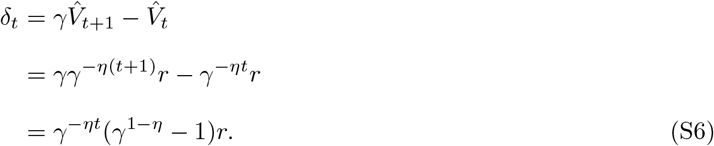

Here, *δ_t+1_* > *δ_t_*. Hence, the RPE should ramp.

#### Ramping Due to State-Dependent Discounting of Estimated Value

Assume the animal underestimates 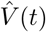 by directly decreasing the temporal discount term *γ*. Then if 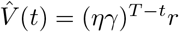, with *η* ∈ (0, 1), we can write in the continuous domain:

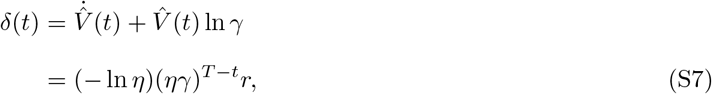

which is monotonically increasing. Hence, the RPE should ramp. Equivalently, in the discrete domain, if 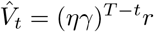 with *η* ∈ (0, 1), we can write

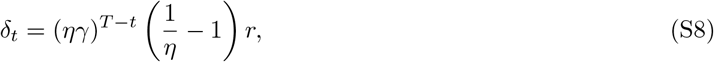

and

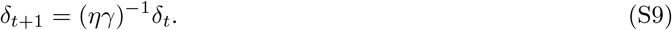

Here, *δ_t_*_+1_ *> δ_t_*. Hence, the RPE should ramp.

### 5 Supplemental Movie 1

Included separately.

